# RanBP2-mediated SUMOylation promotes human DNA polymerase lambda nuclear localization and DNA repair

**DOI:** 10.1101/2020.02.14.949644

**Authors:** M. Moreno-Oñate, A.M. Herrero-Ruiz, M. García-Dominguez, F. Cortés-Ledesma, J.F. Ruiz

## Abstract

Cellular DNA is under constant attack by a wide variety of agents, both endogenous and exogenous. To counteract DNA damage, human cells have a large collection of DNA repair factors. Among them, DNA polymerase lambda (Polλ) stands out for its versatility, as it participates in different DNA repair and damage tolerance pathways in which gap-filling DNA synthesis is required. In this work we show that human Polλ is conjugated with Small Ubiquitin-like MOdifier (SUMO) proteins both *in vitro* and *in vivo*, with Lys27 being the main target of this covalent modification. Polλ SUMOylation takes place in the nuclear pore complex and is mediated by the E3 ligase RanBP2. This post-translational modification promotes Polλ entry into the nucleus, which is required for its recruitment to DNA lesions and stimulated by DNA damage induction. Our work represents an advance in the knowledge of molecular pathways that regulate cellular localization of human Polλ, which are essential to be able to perform its functions during repair of nuclear DNA, and that might constitute an important point for the modulation of its activity in human cells.

## INTRODUCTION

Cellular metabolism daily produces thousands of molecules that are able to generate a variety of DNA lesions, ranging from simple nucleotide changes to DNA breaks [1]. To deal with all DNA lesions, human cells have evolved a variety of repair mechanisms that ensure the maintenance of genome integrity at every cell cycle, avoiding the pathological consequences that might result from a defective response to DNA damage [2, 3]. Short-patch DNA synthesis to fill small gaps is required in virtually all these DNA repair pathways. This gap-filling DNA synthesis is a singular reaction mainly performed by X-family specialized DNA polymerases [4], which develop their abilities in diverse escenarios, such as base excision repair [5], the repair of or tolerance to oxidative damage [6, 7] and the repair of DNA double strand breaks (DSBs) [8–11]. Among the four PolX members identified in human cells to date, namely Polß, terminal deoxynucleotidyl transferase (TdT), Polµ, and Polλ, the latter shows the broadest range of action in DNA repair and tolerance to DNA damage [12]. Functional versatility of Polλ suggests interaction with multiple co-factors in the different pathways in which its gap-filling activity is required, probably implying a complex regulation. In agreement to this, previous studies have revealed cell cycle and DNA damage-dependent phosphorylation-mediated regulation of human Polλ [13–15].

Post-translational modification (PTM) of proteins plays an essential role in the proper functioning and viability of eukaryotic cells. Amongst the different PTMs identified to date, conjugation of Small Ubiquitin-like MOdifier (SUMO) proteins seems crucial in human cells, as it plays relevant roles in key cellular processes, such as DNA replication, transcription and general maintenance of genomic stability [16–19]. SUMOylation is a highly dynamic and transient process that promotes covalent attachment of SUMO moieties to specific lysine residues on target proteins. This reaction is reversible thanks to the existence of a variety of SUMO-specific proteases (SENPs) that efficiently remove SUMO proteins from targeted substrates [16, 17]. Importantly, SUMOylation can protect proteins from ubiquitin-mediated proteasomal degradation, probably competing with ubiquitination machinery for the same target lysine residues [17, 20]. In human cells, there are at least three main SUMO paralogs. SUMO1 shares approximately 45% sequence identity with SUMO2 and 3, and these two are 96% identical to each other. These SUMO proteins are approximately 12 kDa in size and structurally very similar to ubiquitin [20]. SUMO substrates are mainly proteins with nuclear functions, that can be modified either with a single SUMO moiety (mono-SUMO), multiple SUMO units at different sites (multi-SUMO) or with SUMO chains formed at one specific target site (poly-SUMO). In this regard, SUMO paralogs also differ in their ability to form polymeric chains, as SUMO2 and 3 possess internal Lys residues that can serve as SUMO acceptor sites in an auto-modification process that SUMO1 does not have [21]. Likewise, there are substantial differences in the dynamics of SUMO paralogs, because while SUMO1 is mainly found within the nucleoli, the nuclear envelope and cytoplasmic foci, SUMO2/3 are distributed throughout the nucleoplasm [22]. Moreover, whereas SUMO1 mainly exists in the protein-conjugated form, vertebrate cells have a large pool of free SUMO2/3 [22, 23], which is conjugated to high molecular mass proteins when cells are subjected to protein-damaging stimuli [23, 24]. Covalent attachment of SUMO moiety to target proteins occurs in a three-step sequential process, frequently at specific consensus sequences [16, 17]. The SUMO conjugation process is initiated by covalent binding of SUMO protein to the SUMO-activating enzyme SAE1 (E1) through a thioester bond. This is followed by SUMO transference to a second factor, Ubc9 (E2), to form an E2-SUMO1 complex. Although Ubc9 can directly transfer the SUMO to acceptor Lys residues on target protein, this reaction is facilitated *in vivo* by a group of E3 protein ligases, whose mediation is specially relevant for substrate selection in proteins lacking SUMO acceptor consensus sequences [20]. The different SUMO E3 ligases identified so far have distinct subcellular localization, reflecting different biological functions of SUMO modification [25]. Accordingly, whereas RanBP2 is associated with the nuclear pore complex, and has been related to nucleo-cytoplasmic trafficking of specific proteins [26], PIAS, Pc2/CBX4 and others are essentially found in the nucleus, where they play roles in a wide variety of biological processes, from transcription regulation and chromatin remodeling to the maintenance of cellular homeostasis and genome integrity [16,17,20].

Here we report that DNA polymerase lambda (Polλ) is one of the protein targets for SUMOylation in human cells and uncover the biological role of this post-translational modification. We identify human Polλ lysine 27 (K27) as the main acceptor site for the covalent attachment of SUMO proteins both *in vitro* and *in vivo*, and demonstrate that the lack of SUMOylation results in deficient import of Polλ into the nucleus. We also identify the role of nuclear pore complex-associated RanBP2 protein as the main E3 ligase required for Polλ SUMOylation in the cytoplasm, allowing its entry into the nucleus and its function during DNA repair. Overall, our work provides insights into the regulation of Polλ during the DNA damage response and its potential use to control the quantity and/or quality of Polλ-mediated DNA repair events.

## RESULTS and DISCUSSION

### Human Polλ is SUMOylated *in vitro* and *in vivo*

Analysis of human Polλ amino acid sequence by specialized software (SUMOplot, by Abgent; http://www.abgent.com/sumoplot) suggested that Polλ can be modified by SUMOylation. To confirm this prediction, we performed *in vitro* SUMOylation assays using purified proteins (**Figure 1A and Supplemental Figure 1A)**. In these assays we observed the appearance of low electrophoretic mobility species exclusively in the presence of ATP, an essential requirement for SUMO conjugation catalysis. A main modified protein showed an apparent increase in molecular weight of about 15-20 kDa with respect to unmodified protein when either SUMO1 or SUMO2 were included in the assays. This increase was in agreement with mono-SUMOylation of human Polλ (**Figure 1A)**. Moreover, other less represented species with slightly lower electrophoretic mobility were also observed with both SUMO paralogs, suggesting that different lysine residues could be additionally targeted with SUMO, although with less efficiency (**Figure 1A**). We next analyzed whether Polλ SUMOylation also occurred *in vivo* by performing Ni-NTA pulldowns under denaturing conditions from human 293T cells co-expressing Flag-Polλ, His-SUMO1 and Ubc9, the only known E2 conjugation enzyme in mammalian cells. In these experimental conditions, SUMOylation of human Polλ was very efficient, and the size of the different low electrophoretic mobility species observed suggested the formation of poly-SUMO chains (**Supplemental Figure 1B**). Of note, although a fraction of unmodified Polλ was pulled down from all samples, SUMO-Polλ species were stricly dependent on His-SUMO1 over-expression (**Supplemental Figure 1B**). Notably, Polλ SUMOylation was detected at endogenous levels of Ubc9, and was strongly reduced in the presence of a dominant negative Ubc9 mutant that cannot catalyze SUMO conjugation to target proteins [27] (**Supplemental Figure 1B**). SUMOylation of Polλ *in vivo* was also confirmed by using engineered human U2OS cell lines stably expressing His-tagged SUMO paralogs [28, 29]. These cells were transiently transfected with the plasmid encoding for Flag-Polλ, and SUMO conjugates were pulled down from cell extracts as described above. In these experimental conditions, efficient SUMOylation of Polλ was also observed both with SUMO1 and SUMO2 paralogs, being the modification undetectable in control U2OS cells with untagged SUMO proteins (**Figure 1B**). Noteworthy, poly-SUMO chain formation in these experimental conditions was prevented by the molecular features of His-tagged SUMO paralogs expressed in these engineered cells [28, 29].

**Figure 1.**
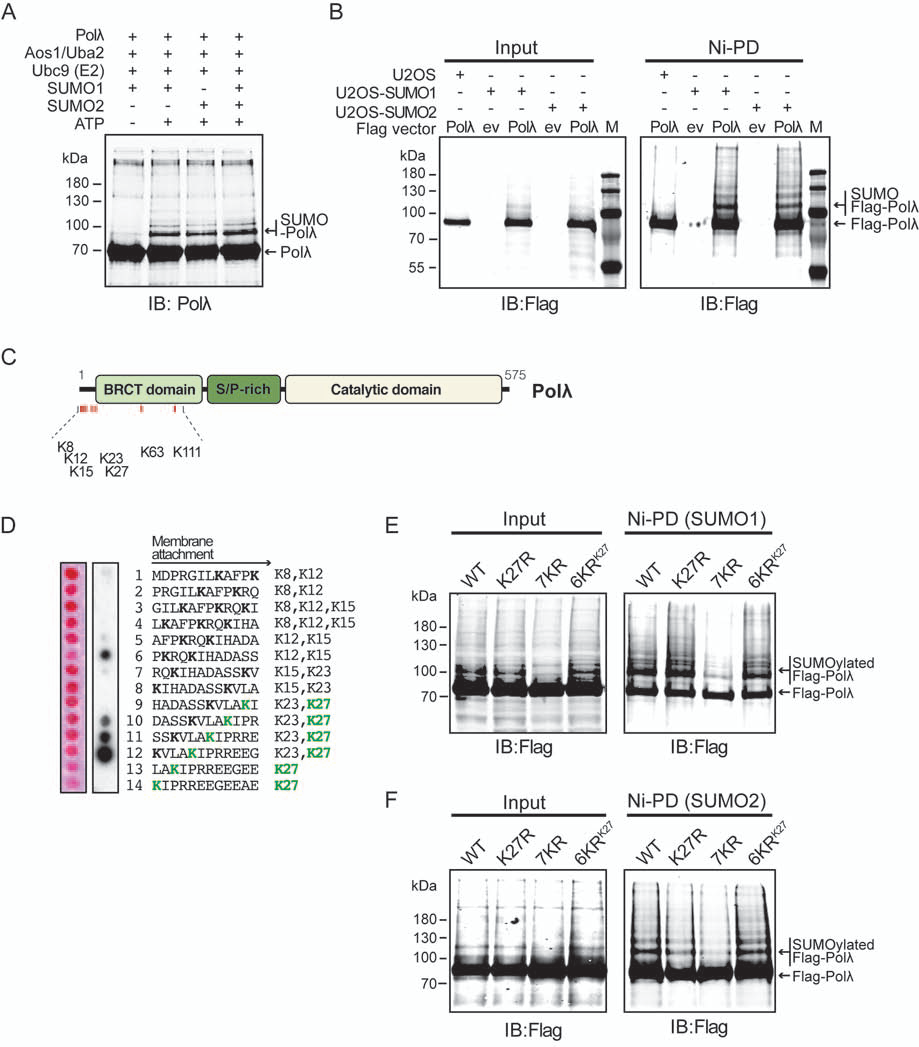
SUMOylation of human Polλ. **(A)** *In vitro* SUMOylation reactions with indicated purified human proteins were carried out as described in Materials and methods. Proteins were resolved by SDS-PAGE and immunoblotted (IB) with anti-Polλ antibody. Unmodified and SUMO-conjugated Polλ are indicated. **(B)** U2OS cells stably expressing either His-SUMO1 or His-SUMO2 were transiently transfected with empty or Flag-Polλ encoding vectors. His-SUMO-conjugates were pulled down on Ni-NTA beads (Ni-PD) under denaturing conditions (Materials and methods) and immunoblotted (IB) with anti-Flag antibody. Expression of Flag-Polλ was monitored by immunoblotting of cell lysates (input). An U2OS cell line without genomic integration of tagged SUMO proteins was assayed in parallel as a control of specificity. Unmodified and SUMO-conjugated Flag-Polλ are indicated. *M* denotes the molecular mass markers in kDa **(C)** Human Polλ domain organization. Main conserved PolX domains [42] and the localization of putative SUMO acceptor lysines in the N-terminal region are indicated. **(D)** Identification of SUMO conjugation sites using peptide arrays. Peptides are numbered sequentially from the starting Met (M) codon. Each spot in the array represents Polλ derived peptides formed by the indicated consecutive residues with 10-amino acid overlap with the previous peptide. Polarity of peptides with respect to membrane attachment is indicated with an arrow. K27 is indicated in green and neighbour lysines are marked in bold. The peptide array was subjected to an *in vitro* SUMOylation assay as described in Materials and methods and then immunoblotted with anti-SUMO1 antibody (right panel). Positive SUMO-conjugated peptides generated dark spots whereas non-conjugated peptides leave blank spots. Left panel represents Ponceau staining of the array. **(E, F)** U2OS cell lines with integrated His-SUMO1 (E) or His-SUMO2 (F) were transiently transfected either with wild-type Flag-Polλ or Flag-Polλ K27R, 7KR and 6KR mutants. His-SUMO-conjugates were pulled down on Ni-NTA beads (Ni-PD) as in (B) and immunoblotted (IB) with anti-Flag antibody. Expression of Flag-Polλ variants was monitored by immunoblotting of cell lysates (input). Unmodified and SUMO-conjugated Flag-Polλ are indicated.

### Identification of Polλ SUMOylation sites

Our initial analysis *in silico* predicted lysine 27 (K27) as the amino acid residue with the highest probability to be targeted by SUMO moiety in human Polλ (**Supplemental Figure 2A and 2B**). K27 is embedded in a highly conserved non-consensus SUMOylation motif located at the most N-terminal region of the protein (A<u>K</u>IP, with the underlined residue being the SUMO target; **Supplemental Figure 2A**). This region is characterized by a great flexibility, that would be in agreement with previously reported SUMO preferential targeting to highly disordered regions [29]. We confirmed initial prediction by performing *in vitro* SUMOylation assays using peptide arrays that covered the most N-terminal region of human Polλ, including both K27 and all its proximal lysine residues (residues 8, 12, 15 and 23; **Figure 1C and 1D**). These assays showed the strongest SUMO targeting in peptides containing K27 (**Figure 1D**), according to proteome-wide studies that had also identified this residue as the main target for Polλ SUMOylation (**Supplemental Figure 2C**) [29]. We therefore generated the most conservative mutation of SUMO acceptor lysine (K27R mutation) and analyzed the effect on SUMOylation *in vivo*, as described above. In these assays, whereas a reduction of SUMO1 conjugation in K27R mutant compared to wild-type Polλ was barely detected, an evident decrease was observed in the case of SUMO2 conjugation (**Figure 1E and 1F**). This effect could be explained to some extent by the lower efficiency of SUMO2 conjugation compared to SUMO1, already observed *in vitro* (**Figure 1B**). Likewise, these assays suggested the existence of alternative SUMOylation target sites in Polλ, also in agreement with results obtained *in vitro* (**Figure 1A**) and with proteomic studies that identified K8 and K23 residues as alternative (secondary) lysines for SUMOylation in the N-terminal region (**Supplemental Figure 2C**) [29]. To prevent SUMOylation of these alternative sites, we generated a mutant in which all seven N-terminal lysine residues, including K8, K12, K15, K23, K27 and two additional and more distant ones (K63 and K111) were equally substituted by arginines (mutant 7KR; **Figure 1C**), and analyzed the effect of these mutations in SUMOylation of Polλ. This analysis showed a dramatic decrease of SUMO conjugation efficiency in 7KR mutant compared with wild-type Polλ, both with SUMO1 and SUMO2 (**Figure 1E and 1F**). We obtained fully confirmation of Polλ K27 residue SUMO-specific targeting *in vivo* through generation of an additional mutant protein in which all N-terminal lysine residues except such K27 were mutated to arginine (mutant 6KR). Notably, this 6KR mutant fully recovered SUMOylation profile observed *in vivo* in wild-type Polλ, both with SUMO1 and SUMO2 (**Figure 1E and 1F**). Altogether, these results indicate that K27 is actually the main target for SUMOylation *in vivo*. It is worth noting that because SUMOylation targets lysine residues of protein substrates, it can potentially compete with other lysine-directed PTMs like acetylation or ubiquitylation [17]. Indeed, Polλ K27 has been identified as a target for ubiquitination, which controls Polλ levels during cell cycle [30]. In light of our results, it is tempting to speculate that K27 SUMOylation could antagonize the effects of ubiquitin, a competitive relationship that has been reported in many other proteins, including some DNA damage response factors [2, 17].

### SUMOylation regulates subcellular localization of human Polλ

One of the first roles assigned to SUMOylation *in vivo* is to regulate subcellular localization of some target proteins, such as RanGAP protein, the first identified SUMO substrate [31]. Therefore, we analyzed by immunofluorescence the subcellular localization of Flag-tagged Polλ versions, including wild-type, mutants analyzed in previous *in vivo* assays (K27R, 7KR and 6KR) and two additional mutants affecting the nearest lysine amino acid residue K23 (K23R and K23/27R) (**Figure 2A**). As expected from a DNA repair polymerase, the distribution of wild-type Polλ in human U2OS cells was predominantly nuclear (more than 92% cells). In contrast, Polλ K27R single mutant showed a markedly different subcellular distribution, with a fraction of the protein remaining outside the nucleus (more than 25%; p<0.01 in Anova test). This effect was specific of K27 amino acid residue, as it was not observed when mutation affected K23 residue (**Figure 2A**). The abnormal distribution seen in K27R mutant was strongly exacerbated in Polλ 7KR, the mutant that was barely SUMOylated *in vivo* (**Figure 1E and 1F**), which was almost completely excluded from the nucleus (**Figure 2A**). Notably, this cytoplasmic distribution was abolished when the K27 residue became available again for SUMOylation, in the 6KR mutant, completely recovering the nuclear localization observed in the wild-type Polλ (**Figure 2A**). These results suggest that SUMOylation at K27 residue controls Polλ entry into the nucleus *in vivo*. Alternatively, SUMO modification might be required to retain Polλ within the nucleus. To evaluate this possibility, we analyzed the effect of leptomycin B (LMB) in the subcellular localization of the Polλ 7KR mutant protein. LMB is an inhibitor of the nuclear-cytoplasmic shuttling of proteins that contain specific nuclear export signals [32], and human Polλ has a predicted nuclear export signal at the BRCT domain. The presence of LMB did not have any effect on the distribution of Polλ K7R protein (**Supplemental Figure 3**), indicating that SUMOylation is not required to retain Polλ inside the nucleus and pointing towards a role in the translocation of Polλ from the cytoplasm into the nucleus. To directly confirm this hypothesis, we generated Polλ SUMOylation mimetic constructs and analyzed their subcellular localization by means of immunofluorescence. These constitutive SUMO-Polλ versions were obtained by translational fusion of mature SUMO1 or SUMO2 to the N-terminus of Polλ K7R mutant protein (**Supplemental Figure 4A**). We also replaced SUMO specific C-terminal double glycine residues by alanine residues in fusion proteins (S1-fusAA and S2-fusAA, respectively) to avoid SUMO removal due to the strong activity of SUMO-specific proteases (SENPs) *in vivo* [16] (**Supplemental Figure 4A**). Genetic fusion of either SUMO1-AA or SUMO2-AA to Polλ 7KR mutant did not restore nuclear localization of Polλ (**Supplemental Figure 4B**), suggesting that the presence of SUMO moiety in the N-terminus of Polλ, although necessary, is not enough to facilitate its entry into the nucleus. Nevertheless, given that both dynamism and reversibility are essential for cellular roles of SUMOylation, it is very likely that these fusion proteins might be incompatible with nucleo-cytoplasmic shuttling functions. A defect in the proper and functional folding of fusion proteins cannot be ruled out either. Regardless, our study uncovers the pivotal role of SUMOylation in the control of cellular distribution of Polλ, and identify the molecular basis of previous observations that Polλ catalytic core, a protein version lacking the amino terminal region including BRTC and Ser-Pro rich domains, is not able to enter into the nucleus [33, 34].

**Figure 2.**
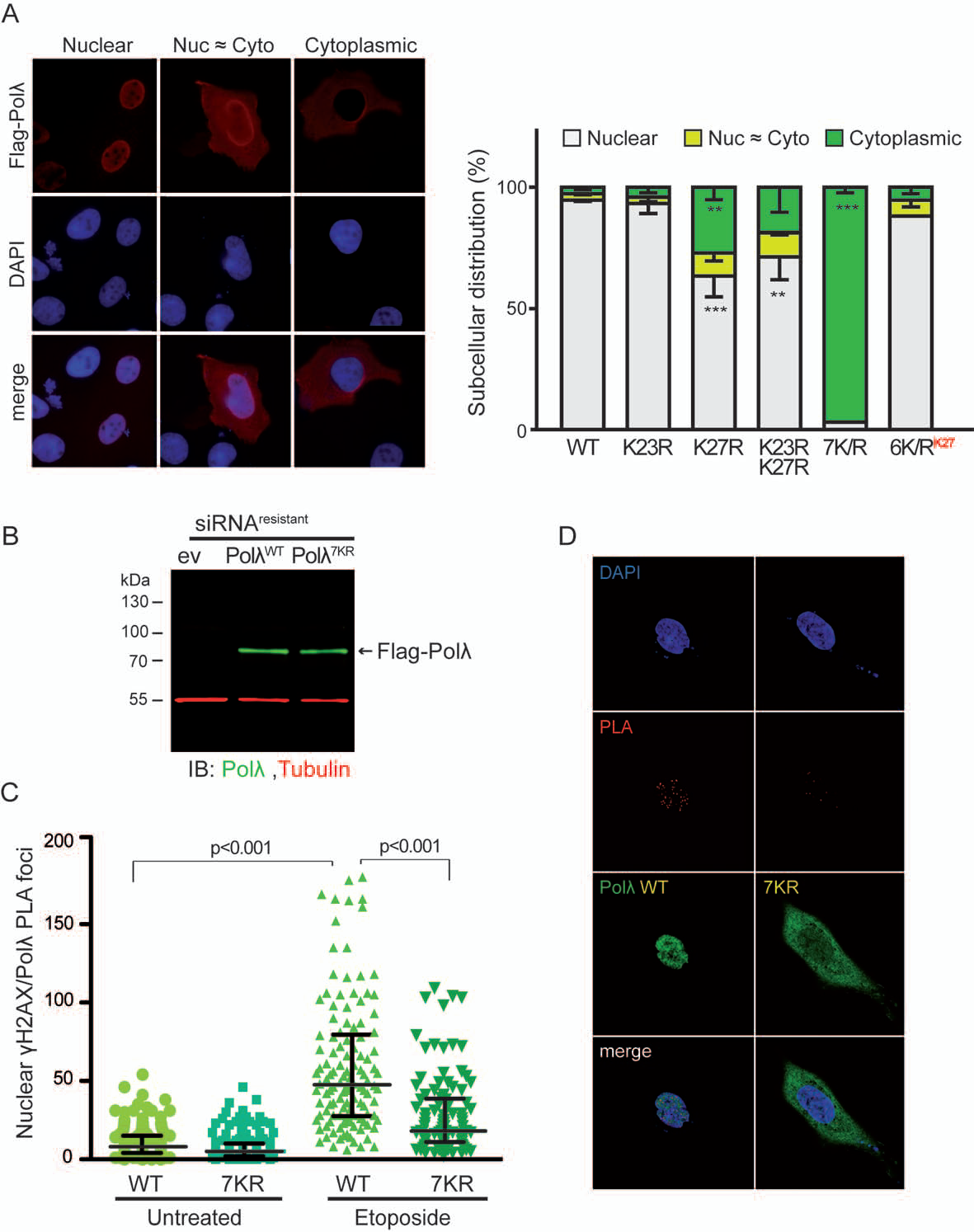
Subcellular localization of Polλ is controlled by SUMOylation. **(A)** Human U2OS cells were transiently transfected with different Flag-Polλ versions (either WT or K23R, K27R, K23/27R, 7KR or 6KR mutants) and subcellular localization of Polλ was examined by immunofluorescence using anti-Flag antibody (red). Nuclei were stained with DAPI (blue). Merge of both Flag and DAPI signals is also shown. Polλ expressing cells were blindly scored for their Polλ subcellular localization, and classified in one of the following three categories: mainly nuclear, equally distributed between nucleus and cytoplasmic (Nuc ≈ Cyto) or mainly cytoplasmic. Representative pictures for each condition are shown in the left panel. The experiment was performed three independent times, with more than 100 independent cells counted each time. The mean percent localization of each condition is plotted, and error bars represent the SD of each data set. Analysis of variance (Anova) was used to test for significance in differences in subcellular localization of Polλ variants (**, p<0.01; *** p<0.001). **(B)** Human His-SUMO1 U2OS cells were treated with specific POLL siRNAs, as previously described [15], and then transiently co-transfected with either Flag-empty vector or siRNA-resistant versions of Flag-Polλ (WT and 7KR). Resistance to siRNA-mediated silencing was confirmed by immunoblotting (IB) with anti-Polλ antibody (green). Immunoblotting with anti-tubulin (red) was used as a loading control. (**C**) Human His-SUMO1 U2OS cells treated as in (B) were subjected or not to etoposide treatment (20 µM, 30 min) and processed for PLA using anti-Polλ and anti-phosphoH2AX (γH2AX) antibodies. PLA foci (red) were counted in transfected cells. Plots display the median values (black bar) for nuclear γH2AX/Polλ PLA foci in each experimental condition (untreated WT, 8.0; untreated 7KR, 5.0; etoposide WT, 47.5; etoposide 7KR, 18.0) plus interquartile range (25th and 75th percentiles). The statistical significance was calculated using GraphPad Prism 5.0; p<0.001 in one-way Anova analysis of variance test of three independent experiments, with >35 individual cells analyzed for each condition in each independent repeat (total number of cells scored: WT untreated, 147; 7KR untreated, 122; WT etoposide, 127; 7KR etoposide, 108). (**D**) Representative images of one experiment described in (C). γH2AX/Polλ PLA foci are shown in red. Flag-Polλ transfected cells (in green) were identified by performing post-PLA staining by using secondary A488-conjugated anti-rabbit antibodies. Nuclei are stained with DAPI (blue).

### Lack of SUMOylation results in decreased Polλ recruitment to etoposide-induced DSBs

Given its relevance in the nuclear localization of Polλ, we wanted to determine the consequences of interfering with SUMOylation on its recruitment to DNA lesions. Among other functions, Polλ is involved in the repair of a subset of DNA DSBs through NHEJ [8,9,15]. To specifically measure Polλ recruitment to DSBs, we developed an experimental approach based on proximity ligation assays (PLA) that uses specific antibodies for human Polλ and phosphorylated histone H2AX (γH2AX), a well known marker of DSBs in mammalian cells [35, 36]. We validated the use of this molecular tool to measure the recruitment of Polλ to specific nuclear DSBs *in vivo* by analyzing the formation of PLA foci of endogenous γH2AX and Polλ in human U2OS cells treated with the DSB-inducing drug etoposide. In these experimental conditions, we observed an increase in the number of PLA foci that was directly proportional to the dose of etoposide used (**Supplemental Figure 5**). Once validated the assay, we performed additional experiments in U2OS cells that were pre-treated with specific siRNAs to remove endogenous Polλ, and that concomitantly overexpressed similar levels of siRNA-resistant versions of either wild-type Polλ or Polλ 7KR mutant (**Figure 2B**). In these experimental conditions, U2OS cells expressing wild-type Polλ showed an increase in Polλ-γH2AX PLA foci in response to etoposide-induced DSBs (untreated WT median 8.0 *vs.* etoposide-treated WT median 47.5; 6-fold increase, p<0.001, Anova test) (**Figure 2C**). Notably, U2OS cells expressing Polλ 7KR mutant showed a decrease in Polλ-γH2AX PLA foci after etoposide-induced DSBs with respect to wild-type Polλ (WT median 47.5 *vs*. 7KR median 18; 3-fold decrease, p<0.001, Anova test) (**Figure 2C and 2D**). These results indicate that recruitment of Polλ to etoposide-induced DSBs is reduced when its SUMOylation-mediated entry into the nucleus is impeded. Noteworthy, some Polλ-γH2AX PLA signal is still observed in cells expressing Polλ 7KR (etoposide 7KR median 18 *vs*. untreated 7KR median 5) (**Figure 2C**), that can probably be due to incomplete silencing of POLL expression. However, alternative routes or complementary mechanisms for Polλ nuclear entry and recruitment to DSBs, independently of the SUMO-mediated pathway, cannot be excluded. Accordingly, novel interactions between Polλ NLS derived peptides and cellular importins have been recently described [37]. Future studies, both structural and functional, will clarify these alternative pathways and putative synergies with the findings uncovered in our study.

### RanBP2 E3 ligase mediates Polλ SUMOylation and nuclear localization

SUMOylation of natural substrates *in vivo* is facilitated by a variety of E3 protein ligases, specially in proteins, as Polλ, lacking consensus SUMO acceptor motifs [16, 25]. To date, the only non-nuclear E3 ligase identified is nuclear pore complex-associated RanBP2 protein, that can form a stable complex with both SUMO1 and Ubc9 throughout the cell cycle and promote the last step in the SUMOylation process [38–40]. Considering this, and the nucleo-cytoplasmic shuttling defect seen in our Polλ mutants (**Figure 2A**), we reasoned that RanBP2 could be the E3 ligase involved in Polλ SUMOylation. Supporting this possibility, proximity ligation assays (PLA) detected physical proximity between RanBP2 and Polλ predominantly in the outer side of nuclear envelope (**Figure 3A and Supplemental Figure 6**). Moreover, we detected direct interaction of these two proteins by co-immunoprecipitation in the absence of any external stimulus (**Figure 3B)**, suggesting that Polλ might be targeted for RanBP2-mediated SUMOylation constitutively. To verify a functional interaction between RanBP2 E3 ligase and Polλ, we performed His-SUMO pulldown assays as those described above in U2OS cells that were previously treated with RanBP2 specific siRNAs. In these experimental conditions, we observed that the reduction in RanBP2 expression led to a strong concomitant decrease in conjugation of SUMO to Polλ (**Figure 3C and 3D**). This effect was not seen in cells treated with control non-targeting siRNAs, so that confirmed RanBP2 as the main E3 ligase involved in Polλ SUMOylation. In the same way, immunofluorescence assays performed in Flag-Polλ expressing U2OS cells that had been silenced for RanBP2 expression also showed a strong decrease of Flag-Polλ nuclear localization when compared to cells treated with control non-targeting siRNAs (**Figure 3E**; more that 40% reduction in nuclear localization; p<0.001 in an unpaired t-test). These results were in fully agreement with those obtained with non-SUMOylatable Polλ mutants in our *in vivo* SUMOylation assays (**Figure 2A**). Altogether, our results indicate that RanBP2 is the main E3 ligase participating in SUMO-mediated modification of human Polλ to promote its nuclear import. Of note, nuclear import of artificial cargos can still occur *in vivo* in RanBP2-depleted cells, although rates are substantially reduced [26]. This could explain residual nuclear Polλ observed in our analyses with K27R single mutant (**Figure 2A**), and, again, suggest the existence of SUMO-independent pathways, still to be deciphered, through which Polλ could enter into the nucleus.

**Figure 3.**
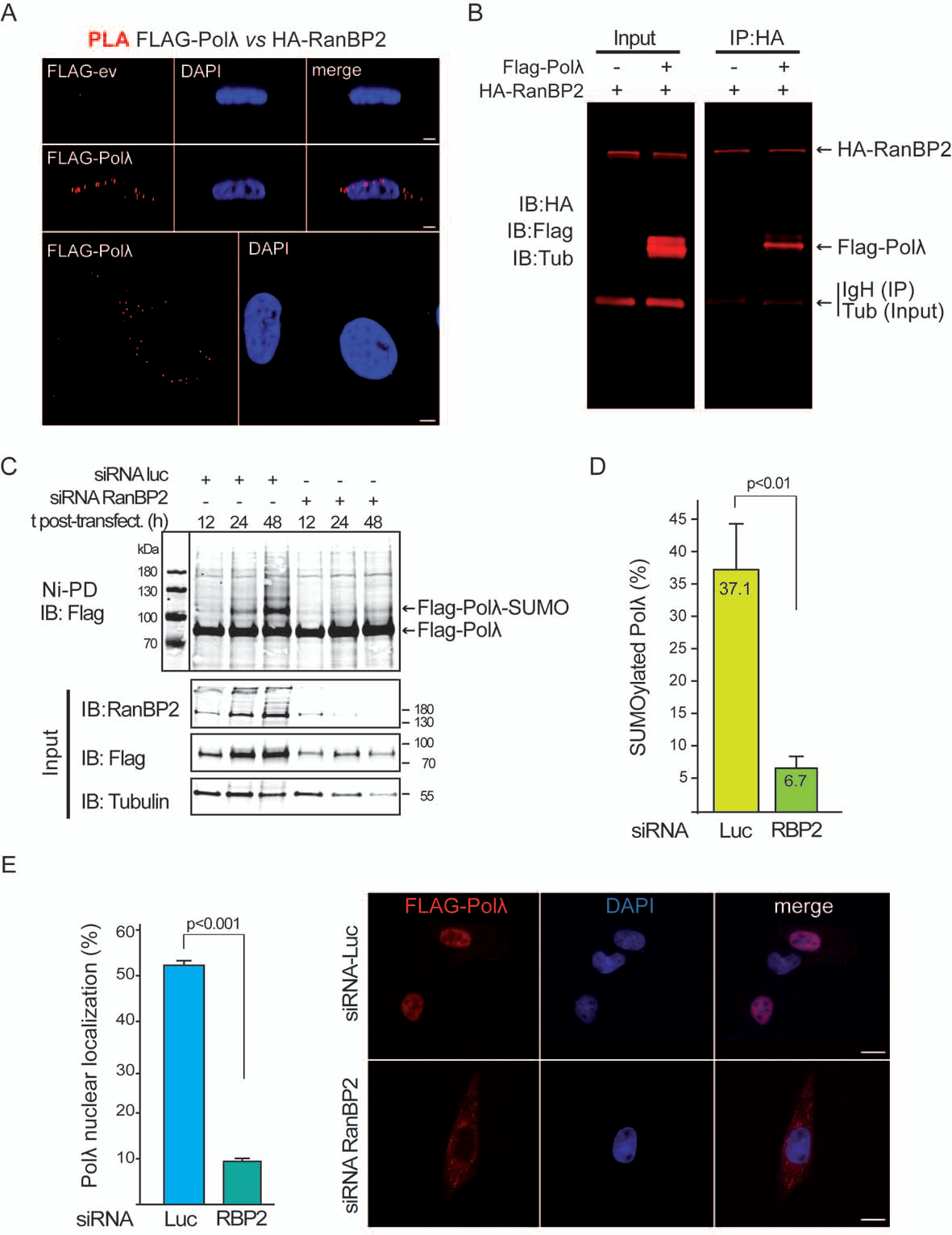
Physical and functional interaction of RanBP2 and Polλ at the nuclear envelope. **(A)** Human U2OS cells were transiently co-transfected with a plasmid encoding for HA-RanBP2 together with either Flag-POLL WT or Flag-empty (ev) vectors. Proximity ligation assays (PLA) were performed by using mouse monoclonal anti-RanBP2 and rabbit anti-Polλ antibodies. Subcellular localization of PLA foci (red) was detected by using confocal laser scanning microscopy and nuclei were stained with DAPI (blue). X/Z views are shown in upper and middle panels, with merged images included in the right panels; an X/Y view is also shown in bottom panels, only with Flag-Polλ WT transfected cells. Scale bars, 10 µm. **(B)** Human 293T cells were transiently transfected as in (A) and HA-RanBP2 was immunoprecipitated 24 h later with anti-HA antibody. Recovered immunocomplexes were analyzed in 4-20% SDS-PAGE and immunoblotting with anti-HA, anti-Flag and anti-tubulin antibodies. **(C)** Human His-SUMO1 U2OS cells were treated either with control (luciferase) or RanBP2 siRNAs and 24 h later transiently transfected with Flag-POLL WT vector. SUMO-Polλ conjugates were pulled down on Ni-NTA beads under denaturing conditions at indicated times post-transfection and immunoblotted with anti-Flag antibody (*upper panel*). Expression of RanBP2, Flag-Polλ and tubulin was monitored by immunoblotting of cell lysates with the corresponding antibodies (input). **(D)** Human His-SUMO1 U2OS cells were treated as in (C) and pulled down on Ni-NTA beads under denaturing conditions performed 48 h after vector transfection. Quantification represents percentage of SUMOylated Polλ in cells (mean value ± standard deviation, SD) for each condition analyzed in three independent experiments. Statistical significance was determined by using an unpaired t-test. **(E)** Human His-SUMO1 U2OS cells were treated as in (C) and subcellular localization of Flag-Polλ was examined by immunofluorescence using anti-Flag antibody (red). Nuclei were stained with DAPI (blue). Representative images of one experiment are shown. Scale bars, 25 µm. Quantification represents percentage of nuclear Polλ in cells (mean value ± standard deviation, SD) for each condition analyzed in three independent experiments, with more than 50 individual cells analyzed for each condition in each independent repeat. Statistical significance was determined by using an unpaired t-test.

### Polλ SUMOylation is enhanced after MMS-mediated DNA damage induction

Once demonstrated the involvement of RanBP2-dependent SUMOylation at the nuclear pore complex in the translocation of Polλ into the nucleoplasm, we wanted to determine if this molecular mechanism was affected by DNA damage. By using engineered human U2OS cell lines stably expressing His-tagged SUMO paralogs described above, we were able to efficiently detect SUMOylation of endogenous Polλ in Ni-NTA pull down assays (**Figure 4A**). This allowed us to evaluate the effect of DNA damage induction on Polλ SUMOylation in more physiological conditions. Such analysis revealed an increase of Polλ SUMOylation when cells were subjected to DNA damage induction, being the increase specially evident in response to the DNA alkylating agent methyl methanesulfonate (MMS) (**Figure 4B and 4C**). MMS-induced DNA lesions are mainly repaired by base excision repair (BER), a repair pathway in which DNA polymerase ß is the main gap-filling DNA polymerase [41, 42]. Although Polλ participation in this repair route was initially suggested as a *back-up* mechanism in mouse cells [43], it has also been shown that cells deficient in both Polλ and Polß are hypersensitive to MMS [5], suggesting that both PolX enzymes synergistically participate in the repair of a common set of DNA lesions. Interestingly, it has been recently reported that human Polλ can also perfectly complement the absence of Polß in POLB-deficient cell extracts, leading authors to suggest that Polλ might be sequestered *in vivo* in a complex with other proteins or post-translationally modified in a way that limits its ability to participate effectively in BER in normal conditions [44]. Overall, our results would be in agreement with a model in which, if Polß is overcome by the amount of damage generated by MMS, a greater requirement of nuclear Polλ would be needed, that might be achieved through the SUMO-mediated translocation of Polλ into the nucleus. It is worth noting that main SUMOylation site in Polλ (K27 residue) is, in turn, target site for ubiquitination, which is a signal for its subsequently degradation via proteasome [30]. Therefore, it cannot be ruled out that SUMOylation at K27 makes Polλ more stable and, therefore, more active in the repair of MMS-induced damage. Future studies will be necessary to validate this model and elucidate the precise role of Polλ SUMOylation in BER regulation. On the other hand, it is equally noteworthy that MMS produces a wide variety of DNA lesions, some of which prevent the progression of replication forks [45]. Therefore, one possibility would be that the increase in Polλ SUMOylation in response to MMS is related to replication forks problems. In agreement with this, replicative stress induced with hydroxyurea (HU) also caused a concomitant increase in SUMOylation of Polλ (**Figure 4C and Supplemental Figure 7**). In spite of this, we did not detect significant differences in Polλ SUMOylation at different stages of a cell cycle in human U2OS cells (**Figure 4D**). Overall, our data suggest that SUMOylation is a relevant signal for human Polλ to entry the nucleus in situations where its gap-filling activity can be specially required, e.g., during MMS-triggered BER or upon replicative stress.

**Figure 4.**
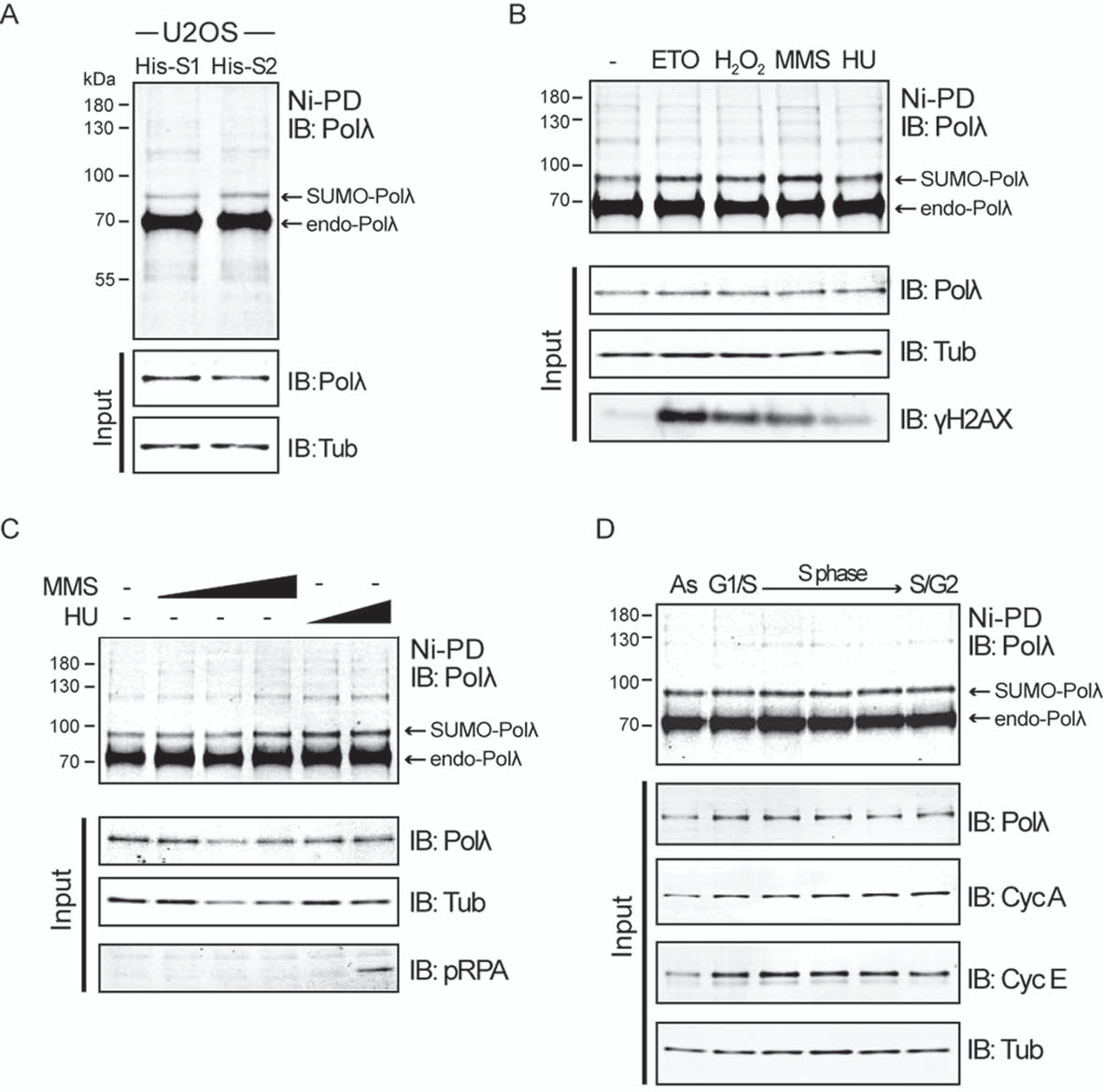
Increased SUMOylation of Polλ in response to DNA damage. **(A)** His-SUMO1 and -SUMO2 U2OS cells were grown in p100 dishes until 80% confluence and then harvested. His-SUMO-conjugates were pulled down on Ni-NTA beads (Ni-PD) under denaturing conditions and endogenous SUMO-Polλ was detected by immunoblotting (IB) using anti-Polλ antibody. Expression of Polλ and tubulin was monitored by immunoblotting cell lysates (input) with the corresponding antibodies. **(B)** His-SUMO2 U2OS cells were grown as described in (A) and either mock-treated or treated with etoposide (20 µM, 30 min), hydrogen peroxide (750 µM, 1 h), MMS (0.03%, 30 min), HU (2 mM, 2h) before harvesting. His-SUMO-conjugates were pulled down on Ni-NTA beads (Ni-PD) and endogenous SUMO-Polλ was detected as described in (A). Expression of Polλ and tubulin was monitored by immunoblotting of cell lysates (input) with the corresponding antibodies. DNA damage induction was confirmed by immunoblotting of cell lysates (input) with anti-phosphorylated H2AX (γH2AX) antibody. (**C**) His-SUMO2 U2OS cells were grown as in (A) and treated with increasing doses of either MMS (0.01%, 0.02% and 0.03%, 30 min) or HU (3 and 5 mM, 2h). His-SUMO-conjugates were detected as described above. Expression of Polλ and tubulin was monitored and replicative stress induction was confirmed with anti-phosphorylated (S4/S8) RPA antibody. (**D**) His-SUMO2 U2OS were synchronized in G1 by using thymidine block, as previously described [56]. Thymidine block was released from G1-enriched cultures and His-SUMO conjugates were pulled down at different times as indicated in (A). Recovering times were selected according to [56]. Cell progression at different stages of the cell cycle was monitored by immunoblotting with anti-cyclin A and E antibodies, and expression of Polλ and tubulin was monitored as described before.

## Concluding Remarks

Detailed knowledge of the post-translational regulation of DNA repair proteins is essential to understand how cells deploy the complicated and tangled network of processes that constitute the DNA damage response. Finely coordinated functioning of this complex network will determine cell viability and prevent the development of age-related diseases and cancer. In particular, it is of great interest to decipher the molecular mechanisms that allow DNA repair factors to enter the nucleus and accomplish their functions, which can have negative consequences for the cell, for example contributing to tumorigenesis as a consequence of inefficient DNA repair [45]. In this work we have identified SUMOylation as a novel post-translational modification of human DNA repair polymerase lambda (Polλ) with a relevant influence in its potential functionality *in vivo*. We have uncovered that SUMO modification fundamentally occurs at N-terminal K27 amino acid residue and requires E3 activity ligase from RanBP2, a nucleoporin present in the outer side of the nuclear envelope. Likewise, we have discovered that nuclear localization of human Polλ is SUMO-dependent, so that such modification is required for Polλ to be recruited to DNA lesions to perform its function. It is worth noting that transport pathways may offer attractive therapeutic targets, as they represent a very selective way of cancelling a biological activity without affecting others. Consequently, the nature of the molecular pathway described here suggests that preventing Polλ SUMOylation by using small molecule inhibitors of this specific modification (for example, blocking SUMO-acceptor sites) could be a suitable point to abolish its polymerase activity in the nucleus, which could be interesting in order to explore the effect of selective Polλ inhibition on tumor cell growth. High levels of replicative stress and excess of oxidative lesions (i.e. ROS) are a hallmark of tumor cells that will demand a great functioning of BER and DNA damage tolerance pathways. Disrupting the cellular mechanisms that cope with this aberrant situation, for example by inhibiting the ATR or Chk1 kinases, constitutes a very promising strategy for cancer treatment [47, 48]. Interestingly, the suppression of Polλ activity induces synthetic lethality when combined with Chk1 inhibitors [49], what turns Polλ and its nucleo-cytoplasmic shuttling as additional potential chemotherapeutic target to be analyzed in more detail. Importantly, targeting the N-terminal region of Polλ would solve undesired effects of many of Polλ inhibitors identified so far, that can also affect other polymerases, in particular those from PolX family, due to high similarity in their catalytic center [50, 51]. Although further structural studies should be needed to gain molecular insights of Polλ N-terminal region, some pioneering work performed with the murine ortholog already shed some light in this regard [33]. Finally, another attractive scenario to be explored regarding Polλ SUMOylation inhibition would be related with CRISPR-Cas based gene editing tools. These systems strongly rely on the processing of directed DSBs by the cellular NHEJ repair machinery. The possibility of controlling CRISPR-based gene editing accuracy by pharmacological modulation of the repair process is being intensively evaluated at present [52]. In this regard, the search for selective inhibitors against Polλ SUMOylation could also be useful in the gene editing toolkit, given that a large subset of breaks generated by CRISPR-Cas systems requires specialized DNA gap-filling activity by PolX polymerases as Polλ [53, 54].

## MATERIALS AND METHODS

### Cell cultures

Human embryonic kidney 293T cells and human osteosarcoma cells (U2OS, U2OS HIS-SUMO1 and -SUMO2 expressing cells) [28, 29] were cultured in DMEM medium (Sigma) supplemented with 10% fetal bovine serum (Sigma), 2 mM L-glutamine and antibiotics (100 units/ml penicillin, 100 µg/ml streptomycin; Sigma) at 37°C in a humidified atmosphere containing 5% CO2. Transient transfections of plasmids were performed with Lipofectamine (Life Technologies) according to manufacturer’s instructions.

### Plasmid constructs and siRNA

Wild-type POLL cDNA was amplified by PCR and cloned in the p3xFlag-Myc-CMV expression vector (Sigma) as previously described [15]. POLL mutants were generated by site directed mutagenesis using p3xFlag-[POLL]-Myc-CMV and overlap extension PCR methodology with oligonucleotides listed in Table S1. POLL K6R mutant was generated by site directed mutagenesis that reverted K27R mutation from p3xFlag-[POLL 7KR]-Myc-CMV plasmid, and both Polλ and Polλ7KR siRNA resistant clones were generated by using the oligonucleotides indicated in Table S1 using the p3xFlag-[POLL]-Myc-CMV and p3xFlag-[POLL 7KR]-Myc-CMV plasmids, respectively. All mutated cDNAs were verified by DNA sequencing. Constitutive SUMO-Polλ versions (both with SUMO1 and SUMO2) were generated by PCR amplification of either SUMO1 or SUMO2 cDNA by using oligonucleotide primers with flanking *Bgl*II cut sites (see Table S1). PCR products were digested with *Bgl*II, cloned into p3xFlag-[POLL]-Myc-CMV expression vector and sequenced to verify proper orientation of fusion cDNAs. Site directed mutagenesis of the C-terminal di-glycine motif of SUMO proteins was performed on these plasmids by using oligonucleotides listed in Table S1 and overlap extension PCR methodology. Efficient silencing of endogenous POLL and RANBP2 genes was achieved by two consecutive rounds of transfection of previously validated double-stranded siRNAs (Table S1) [15] into human U2OS cells. Once seeded, cells were cultured for 24 hours and then transfected either with control (luciferase), POLL or RanBP2 specific siRNAs by using RNAiMAX (Life Technologies) following manufacturer’s instructions. Twenty-four hours later, cells were subjected to a second siRNA transfection with the same siRNAs and, when indicated, either p3xFlag-[POLL]-Myc-CMV expression vector or the corresponding control empty vector using Lipofectamine2000 (Life Technologies) according to manufacturer’s instructions. Silencing was confirmed by Western blotting analysis. The siRNA resistant wild-type Flag-POLL was previously described [15], and siRNA resistant clones of Flag-POLL 7KR were obtained by using oligonucleotides listed in Table S1 and Q5 Site Directed Mutagenesis kit (NEB) following manufacturer’s instructions.

### Proteins and *in vitro* SUMOylation assays

Mouse Ubc9, Aos1 and Uba2 were produced in *E. coli* DH5α at 20°C as GST fusions and purified with Glutathione Sepharose 4B beads (GE, Healthcare) according to manufacturer’s instructions. In the case of Ubc9, GST moiety was excised by using the PreScission protease (GE, Healthcare). Human DNA polymerase lambda (Polλ) was a gift of Dr. Luis Blanco (CBM-SO, Madrid, Spain). *In vitro* SUMOylation assays with murine SUMO proteins were performed with 300 ng of purified Polλ. The reaction was carried out in 20 µl of standard SUMO reaction buffer (20 mM HEPES pH 7.5, 50 mM NaCl, 4 mM MgCl2, 0.05% Tween and 1 mM DTT buffer) containing 200 ng Aos1/Uba2 mix (E1), 600 ng Ubc9 (E2) and 250 µM of the corresponding SUMO protein. Reactions were initiated with 250 µM ATP, incubated at 30°C for 3 h and stopped with ß-mercaptoethanol containing Laemmli buffer. Reactions with human SUMO proteins were performed using 500 ng of purified human Polλ and commercially available *in vitro* SUMOylation kits (Boston Biochemicals), following manufacturer’s instructions. After *in vitro* reactions, proteins were run in SDS-PAGE 10% and immunoblotted by using anti-Polλ antibody (A301-640A Bethyl).

### In vitro SUMOylation on peptide arrays

For *in vitro* SUMOylation assays on peptide-scanning arrays, N-terminal acetylated overlapping dodecapeptides covering the N-terminal region of human Polλ were generated by automated spot synthesis onto an amino-derivatized cellulose membrane (CNB Proteomics Core Facility, Proteored, Spain). Peptides were immobilized by their C-termini via a polyethylene glycol spacer. Overlapping peptides were spotted onto membrane so that they shared 10 amino acids with its adjacent peptide on the array, corresponding to a change of two amino acids per peptide. Peptide array membrane was blocked overnight in 1% BSA and 3% Tween-20 before SUMO-conjugation on cellulose-bound peptides. Briefly, enzymatic reaction was performed in a reaction mixture containing 5mM ATP, 50mM NaCl, 5mM MgCl2, 0.2M DTT, 1% BSA and 3% Tween-20, and 0.15 µM E1, 0.20 µM E2 and 0.4 µM SUMO1 proteins for 30 min at 37°C. Nonspecifically bound proteins were washed off by sonication for 2 min in a bath containing 1% SDS, 0.1% ß-mercaptoethanol and 100mM Na2HPO4. Further nonspecific binding was blocked by incubation with 3% BSA-1% Tween-20 in TBS for 1 hour. Subsequently, the membrane was incubated for 2 hours with anti-SUMO1 primary antibody diluted in blocking solution (1:1,000 dilution). Membranes were washed with TBS-Tween-20 (0.1%) and subsequently incubated with HRP-conjugated secondary antibody diluted in blocking solution for 1 hour. Membranes were washed extensively with TBS-T (0.1%) and SUMO1 signal was detected by ECL incubation following the manufacturer’s instructions.

### Immunoblotting, immunoprecipitation and pulldown assays

For western blotting analyses, 2.5x10^5^ human 293T cells were seeded per well in 6-well dishes, cultured for 24 h and transfected with the corresponding plasmids/siRNAs using Lipofectamine 2000. Cells were washed with cold-phosphate buffered saline (PBS) and collected by low speed centrifugation. Cells were then lysed in 500 µL lysis buffer (20 mM Tris-HCl pH 7.5, 150 mM NaCl, 10% glycerol, 2 mM EDTA, 1% NP-40, 1 mM phenylmethylsulfonyl sulfate (PMSF), protease inhibitor cocktails (Sigma) and 1 mM DTT), homogenized and incubated on ice for 30 min. Lysates were cleared by centrifugation at 13,000 rpm for 20 min at 4°C and proteins were resolved by 10-12% SDS-PAGE. For RanBP2 detection proteins were resolved by 4-20% gradient SDS-PAGE gels (Biorad). In any case, proteins were transferred onto Immobilon-FL PVDF membranes (Millipore) by using a wet-transfer system (Biorad) at 150 mA and 4°C during 2 hours. Immunoblotting was performed according to Odyssey LI-COR Biosciences instructions. The following primary antibodies were used for immunoblotting analysis: mouse monoclonal anti-Flag (M2, Sigma), mouse anti-tubulin (T9026, Sigma), rabbit anti-Polλ (A301-640A, Bethyl) and mouse monoclonal anti-RanBP2 antibody (sc-74518, Santa Cruz). After washes with TBS-0.1% Tween20, membranes were incubated with the corresponding IRDye 800CW/ 680RD secondary antibodies (LI-COR Biosciences; 1/15,000) supplemented with 0.1% Tween20, washed again and were allowed to dry to be then analysed in Odyssey infrared imaging system with the ImageStudio Odyssey CLx Software (LI-COR). Quantification of relative band intensities was carried out using ImageJ software. For immunoprecipitations, 293T cells were cultured as before and co-transfected with both p3xFlag-POLL-Myc-CMV and HA-RanBP2 plasmids using Lipofectamine 2000 according to the manufacturer’s instructions. After 24 h, cells were harvested, lysed in 500 µL lysis buffer and incubated on ice for 30 min. Lysates were cleared by centrifugation as indicated before and input samples saved for subsequent analysis. Remaining supernatants were incubated with mouse monoclonal anti-HA antibody (Sigma) overnight at 4°C. Immuno-complexes were then incubated with Protein G-coupled Dynabeads (Life Technologies) for 4 h with end-to-end mixing at 4°C. Beads were washed twice in lysis buffer and bound immuno-complexes were released by boiling samples in Laemmli buffer. Proteins were resolved by 4-20% gradient SDS-PAGE gels (Biorad) and processed for western blotting as indicated before. Input lysate (10%) was loaded alongside unless otherwise stated. For His-tagged SUMO pulldown assays, 3x10^5^ human U2OS HIS-SUMO1- and -SUMO2 expressing cells were seeded in P60 dishes, cultured for 24 h and then transfected with 3xFlag-POLL-Myc-CMV expression vectors as indicated before. After 48 h, cells were washed and collected by centrifugation in ice-cold PBS and stored until processing. For pulldown assays to detect endogenous SUMOylation of Polλ, cells were seeded in P100 dishes and 2-4x10^6^ cells were processed. In any case, cell extracts were then prepared under denaturing conditions by incubation in 500 µL of urea lysis buffer (8 M urea, 0.1 M Tris/HCl pH 6.8, 0.2% Triton-X100) during 30 min with rotation at room temperature. Lysates were cleared by centrifugation at 13,000 rpm for 10 min at room temperature. Input samples were saved and His-tagged SUMO-conjugated proteins were pulled down from cell extracts by using Ni-NTA agarose beads (Life Technologies). Pulled down proteins were washed 3 times in lysis buffer supplemented with 10 mM imidazole and finally eluted in lysis buffer supplemented with 250 mM imidazole and Laemmli buffer. Eluted SUMO-conjugated proteins were boiled and loaded onto 10% SDS-PAGE. Input lysate (5%) was loaded alongside unless otherwise stated. When indicated, cells were treated with different drugs and DNA damaging agents: etoposide, hydrogen peroxide, MMS and HU (Sigma). After treatment, cells were rinsed using PBS and processed as indicated before.

### Immunofluorescence

For standard immunofluorescence studies approximately 2×10^4^ U2OS cells were seeded onto sterile glass coverslips in each well of a four-well dish, cultured for 24 hours and transfected with Flag-POLL constructs as described above. For immunofluorescence studies involving siRNA-mediated silencing of endogenous Polλ or RanBP2, 2.5×10^4^, U2OS cells were seeded onto sterile glass coverslips in each well of a four-well dish and cultured for 24 hours. Silencing of endogenous POLL or RANBP2 was performed as described above. After 72 h, cells were fixed by treatment with 4% formaldehyde in PBS at room temperature for 10 min, and then permeabilized by treatment with 0.2% Triton X-100 in PBS. Cells were incubated with PBS-1% BSA for 30 min to block non-specific antigens, and then were incubated with mouse monoclonal anti-Flag M2 (Sigma; 1/5,000) diluted in PBS-1% BSA. After washes with PBS-0.1% Tween20, cells were incubated with Alexa 594 Fluor-conjugated secondary antibodies (Jackson; 1/1,000) and washed again as described above. Finally, they were counterstained with 4,6 diamidino-2-phenylindole (DAPI 1 µg/ml; Sigma) and mounted using Vectashield mounting medium (Vector Laboratories). Samples were visualized and pictures taken by using a Leyca DM6000B fluorescence microscope with a HCX PL APO 63x/NA 1.40 oil immersion objective. For proximity ligation assays (PLA) human U2OS cells were grown on sterile glass coverslips and co-transfected with both HA-RanBP2 and Flag-Polλ as described before. For PLA assays to analyze and measure endogenous Polλ and γH2AX proximity, transfection step was omitted. After 48 hours, cells were fixed and permeabilized as described above. PLA assays were performed using the Duolink PLA kit (Sigma) according to manufacturer’s instructions with following antibodies: rabbit anti-Polλ (A301-640A Bethyl; 1/1,000), mouse monoclonal anti-RanBP2 antibody (sc-74518, Santa Cruz; 1/1,000) and mouse monoclonal anti-γH2AX (clone JBW301, Millipore; 1/1,000). For PLA assays measuring transfected Flag-Polλ (either si-RNA resistant WT or 7KR mutant) and γH2AX proximity, positively transfected cells were identified by performing post-PLA staining by using Alexa A488 fluor-conjugated anti-rabbit secondary antibodies (Jackson; 1/1,000). After final wash step, cells were counterstained with DAPI and mounted as described above. Confocal analysis was performed with a Leica confocal microscope TCS SP5, using a HCX PL APO lambda blue 63x 1.4 objective with zoom 3. Image stacks were captured keeping a step size of 0.5 microns and sequential scanning was defined for each channel using 543 nm, 488 nm and 405 nm laser lines, respectively.

## ACKNOWLEDGEMENTS

A. Vertegaal (Leiden University Medical Center, The Netherlands) for U2OS expressing His-SUMO1 and -2 cell lines; L. Blanco (CBM-SO CSIC/UAM, Spain) for purified human Polλ.

## AUTHOR CONTRIBUTIONS

JFR conceived and designed the research; MMO, AHR, MGD, and JFR performed the experiments, collected and analyzed; FCL provided scientific input and designed the research; JFR wrote the manuscript with input from all authors.

## TABLE AND FIGURE LEGENDS

**Supplemental Figure 1.**
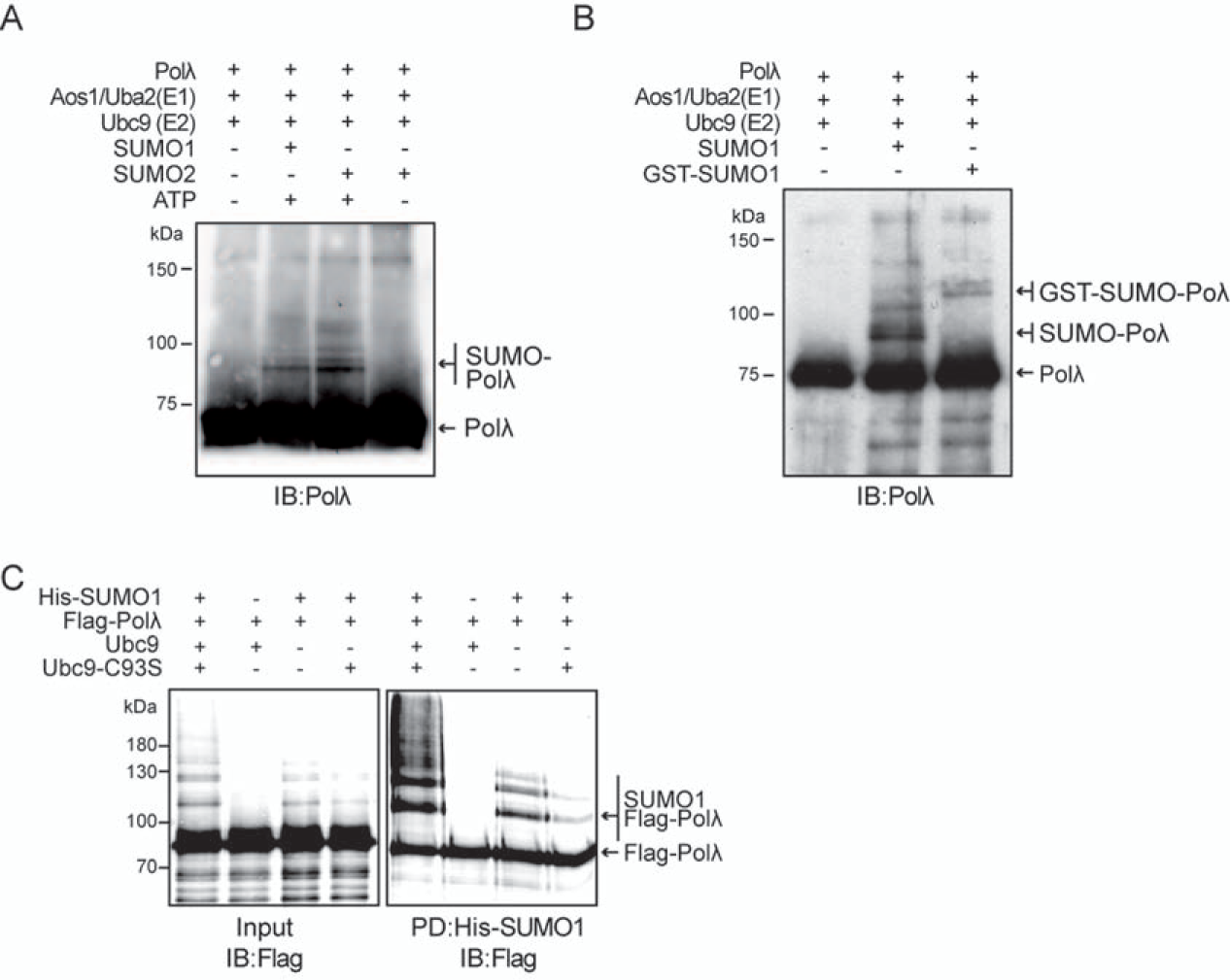
SUMOylation of human Polλ **(A)** *In vitro* SUMOylation assay. Reactions were carried out with purified human Polλ and the indicated purified murine SUMOylation proteins in standard SUMO reaction buffer (see Materials and methods). When indicated, ATP was added to trigger catalytic reactions leading to SUMO conjugation. Proteins were resolved by SDS-PAGE and Polλ was detected by immunoblotting (IB) with anti-Polλ antibodies. *In vitro* reactions produced ATP-dependent Polλ-SUMO conjugates with slower electrophoretic migration compared with unmodified Polλ. Both unmodified and SUMO-conjugated Polλ are indicated. **(B)** *In vitro* SUMOylation assays were performed as in (A), including a GST-SUMO1 fusion protein to confirm Polλ SUMOylation. Proteins were resolved by SDS-PAGE and Polλ was detected by immunoblotting (IB) with anti-Polλ antibodies. The corresponding Polλ-SUMO conjugates with slower electrophoretic migration compared with unmodified Polλ obtained are indicated. GST-SUMO1-Polλ species are in agreement with additional 25 kDa size corresponding to GST protein. **(C)** *In vivo* SUMOylation assays. Human 293T cells were transiently transfected with plasmids encoding for Flag-Polλ, His-SUMO1 and Ubc9. SUMO conjugates were purified from cell lysates 48 h later by Ni-NTA pulldown (PD) assays under denaturing conditions. Polλ SUMOylation was detected by immunoblotting (IB) using anti-Flag antibodies. SUMO-Polλ conjugates are observed as slower migrating species compared with unmodified Flag-Polλ. Both unmodified and SUMO-conjugated Polλ are indicated.

**Supplemental Figure 2.**
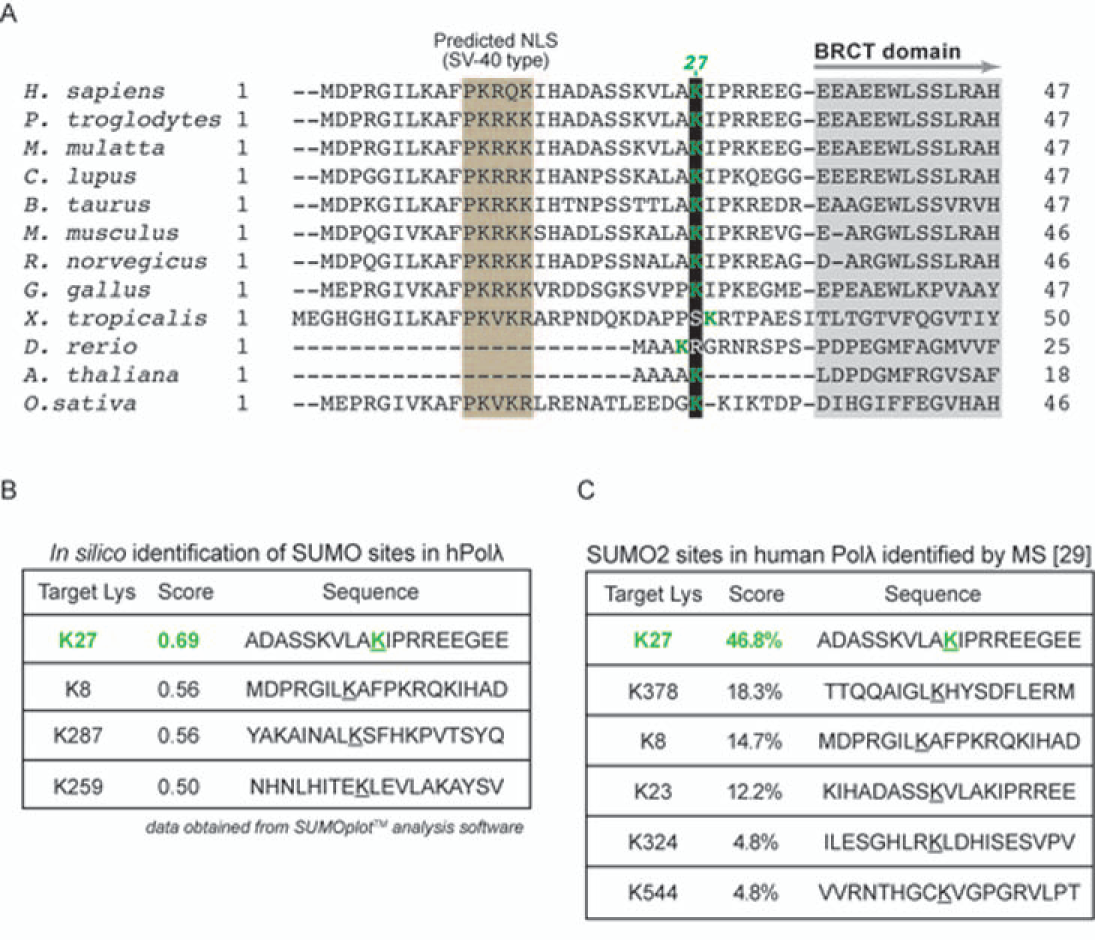
Identification of SUMO sites in human Polλ. (**A**) Sequence alignment of Polλ N-terminal regions from different species. Lysine 27 (K27) is embedded into the non-consensus site for SUMOylation (AKIP in human sequence), and is marked in green over black background. An acidic patch located downstream from K27 is marked in red. The beginning of the BRCT domain (amino acid residues 35-125) and a predicted nuclear localization signal (amino acid residues 11-17) are also indicated [55]. (**B**) Identification of Polλ SUMO conjugation sites *in silico*. Analysis of human Polλ amino acid sequence with SUMOplot^TM^ software identifies lysine 27 as the residue with the greatest potential to be modified by SUMO conjugation. Only lysine residues with highest score are shown. (**C**) Identification of SUMO2 conjugation sites in human Polλ by proteome-wide assays [29].

**Supplemental Figure 3.**
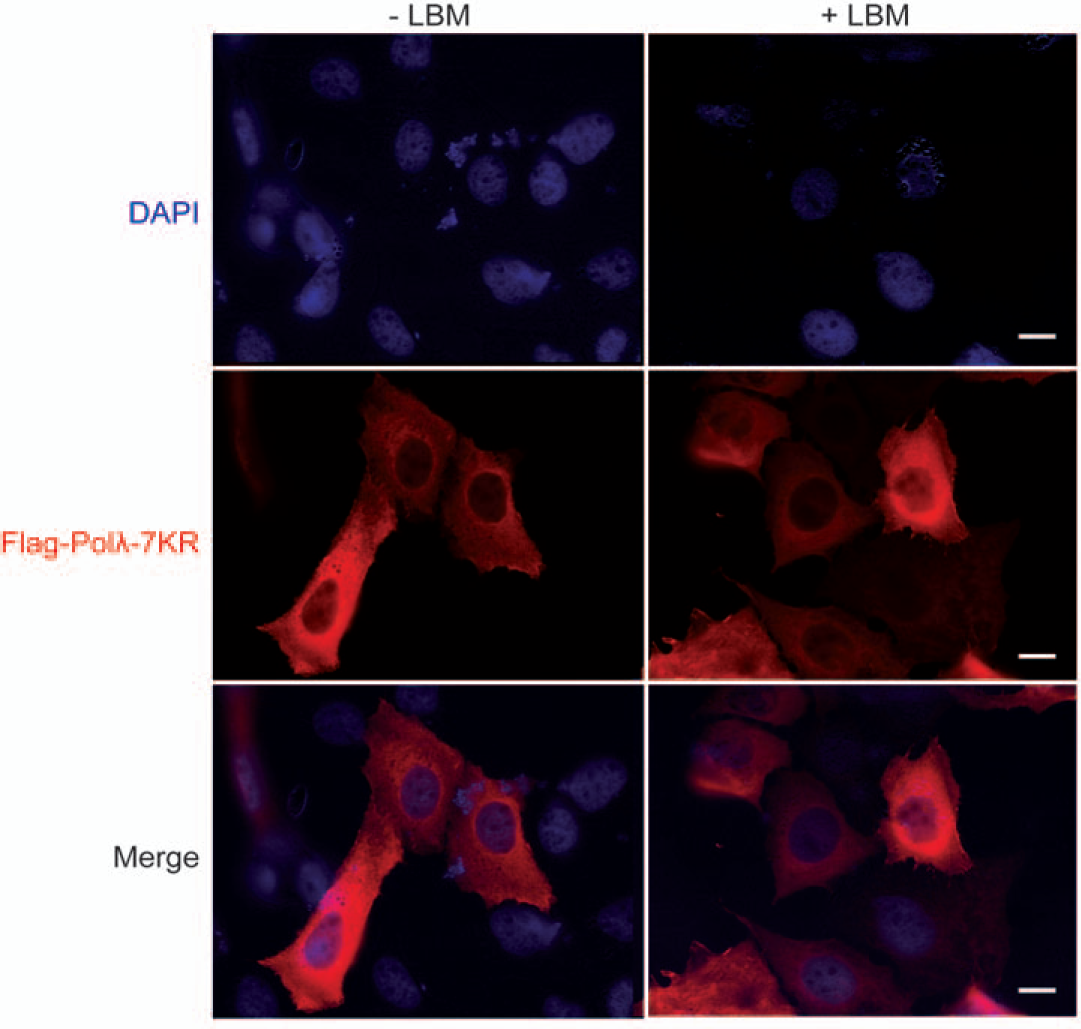
Effect of leptomycin B in Polλ 7KR subcellular localization. Human U2OS cells were transiently transfected with Flag-tagged Polλ 7KR mutant. After 24 h, transfected cells left untreated (-LMB) or treated with 10 ng/µl LMB for 6   h (+LMB), and cellular localization of Flag-Polλ was examined by immunofluorescence staining using anti-Flag antibody (red). Nuclei were stained with DAPI (blue). Merge of both Flag and DAPI signals is also shown. Scale bars (bottom-right corner), 10 µm.

**Supplemental Figure 4.**
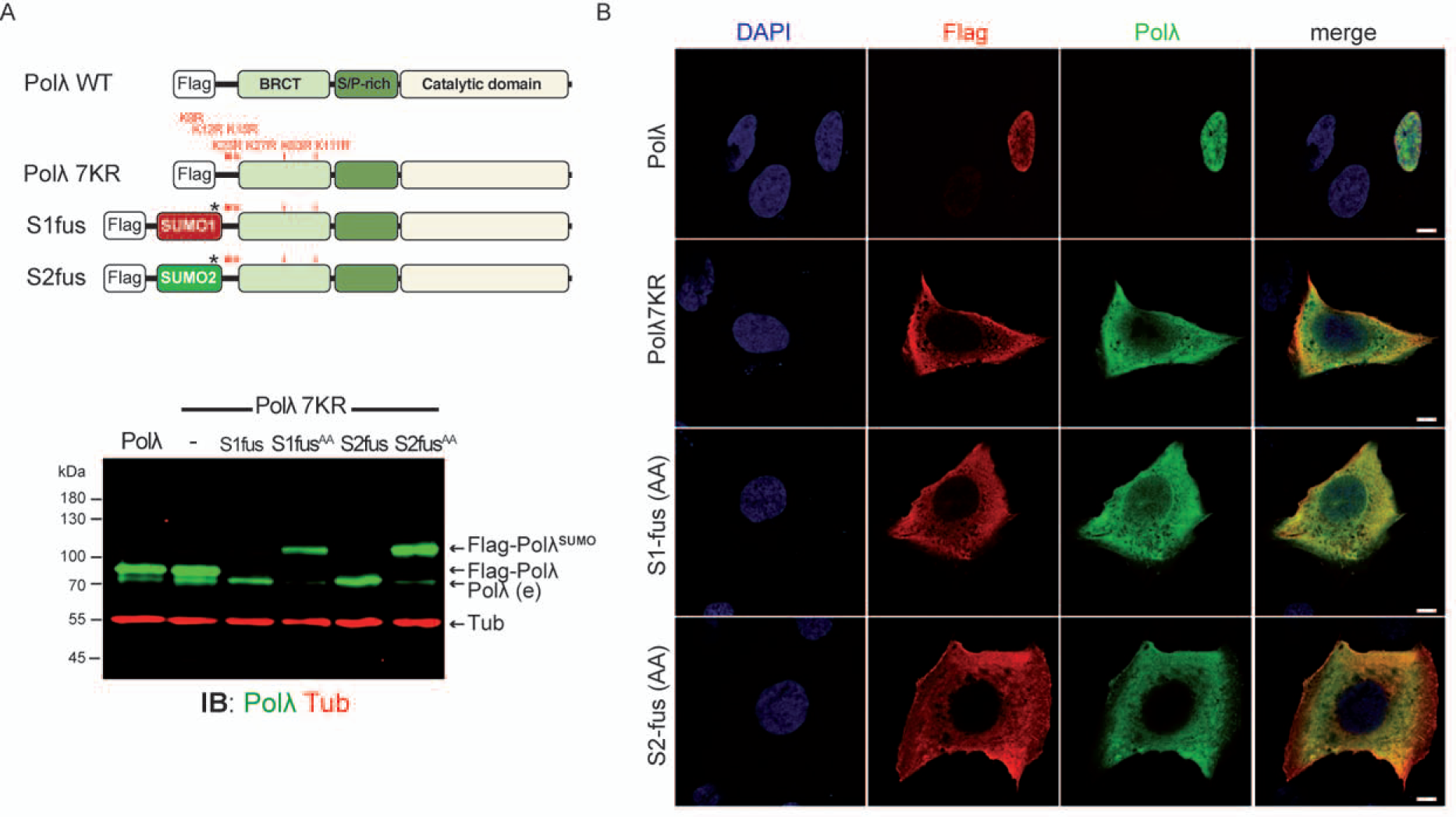
Subcellular localization of constitutive SUMO-Polλ versions (**A**) (*Upper*) Scheme of constitutive SUMO-Polλ 7KR constructs generated by the in-frame fusion of SUMO1 or SUMO2 human cDNA at the N-terminus of Polλ cDNA in the 3xFlag-POLL7KR-myc mammalian expression vector to generate S1-fus and S2-fus SUMO-Polλ7KR fusion proteins, respectively. Localization of the C-terminal di-glycine (GG) motif critical for SUMO conjugation is marked with an asterisk. GG motif was mutated to di-alanine (AA) in the mutants S1-fus and S2-fus to impede SENP proteases action that cause breakage of SUMO-Polλ conjugation (*bottom panel*). Human Polλ conserved PolX domains are: BRCT, BRCA1 C-terminal domain; SP-rich, Ser-Pro rich domain; catalytic, Polß-core domain [56]. The localization of lysine residues in the N-terminal region of Polλ is indicated. (*Bottom*) Human U2OS cells were transiently transfected with vectors encoding the indicated Flag-tagged Polλ versions and expression of corresponding fusion proteins was analyzed by immunoblotting of whole cell extracts with anti-Polλ antibody (green). Tubulin (red) was used as a loading control. In-frame SUMO-Polλ fusions S1-fus (AA) and S2-fus (AA) generated proteins with increased molecular weight over Flag-tagged Polλ according to SUMO moiety (around 20 kDa). S1-fus (GG) and S2-fus (GG) proteins were degraded by SENPs leading to an overexpressed product with the same size as endogenous Polλ, also indicated. (**B**) Subcellular localization of Flag-tagged Polλ wild-type and 7KR mutants, including constitutive SUMO-Polλ fusion proteins S1-fus (AA) and S2-fus (AA), using confocal laser scanning microscopy. U2OS cells were transiently transfected with the indicated plasmids and cellular localization of Flag-Polλ was examined by immunofluorescence using both anti-Flag (red) and Polλ (green) antibodies. Nuclei were stained with DAPI (blue). Representative pictures for each condition are shown. Scale bars, 7.5 µm.

**Supplemental Figure 5.**
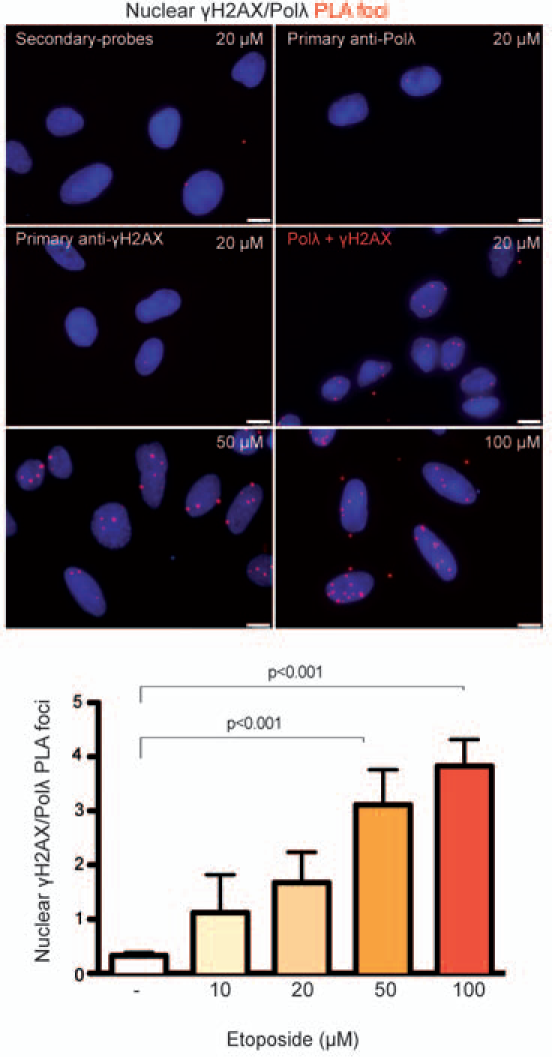
Proximity ligation assays to measure Polλ-mediated DSB repair. Human U2OS cells were mock treated or exposed to the indicated doses of etoposide for 30 minutes to induce DNA DSBs. Cells were then processed for proximity ligation assays (PLA) using rabbit anti-Polλ and mouse anti-phosphorylated H2AX (γH2AX) antibodies. Cells were blindly scored for PLA foci (in red) and nuclei were stained with DAPI (blue). Plot displays the mean values (± standard deviation, SD) of three independent experiments, with more than 50 individual cells analyzed for each condition in each independent repeat. Total number of independent cells scored: untreated, 175; 10 µM, 346; 20 µM, 284; 50 µM, 289; 100 µM, 303. The statistical significance was calculated using GraphPad Prism 5.0; p<0.001 in one-way Anova analysis of variance test. Representative images of one experiment are shown in the upper panel. Scale bar represents 10  µm.

**Supplemental Figure 6.**
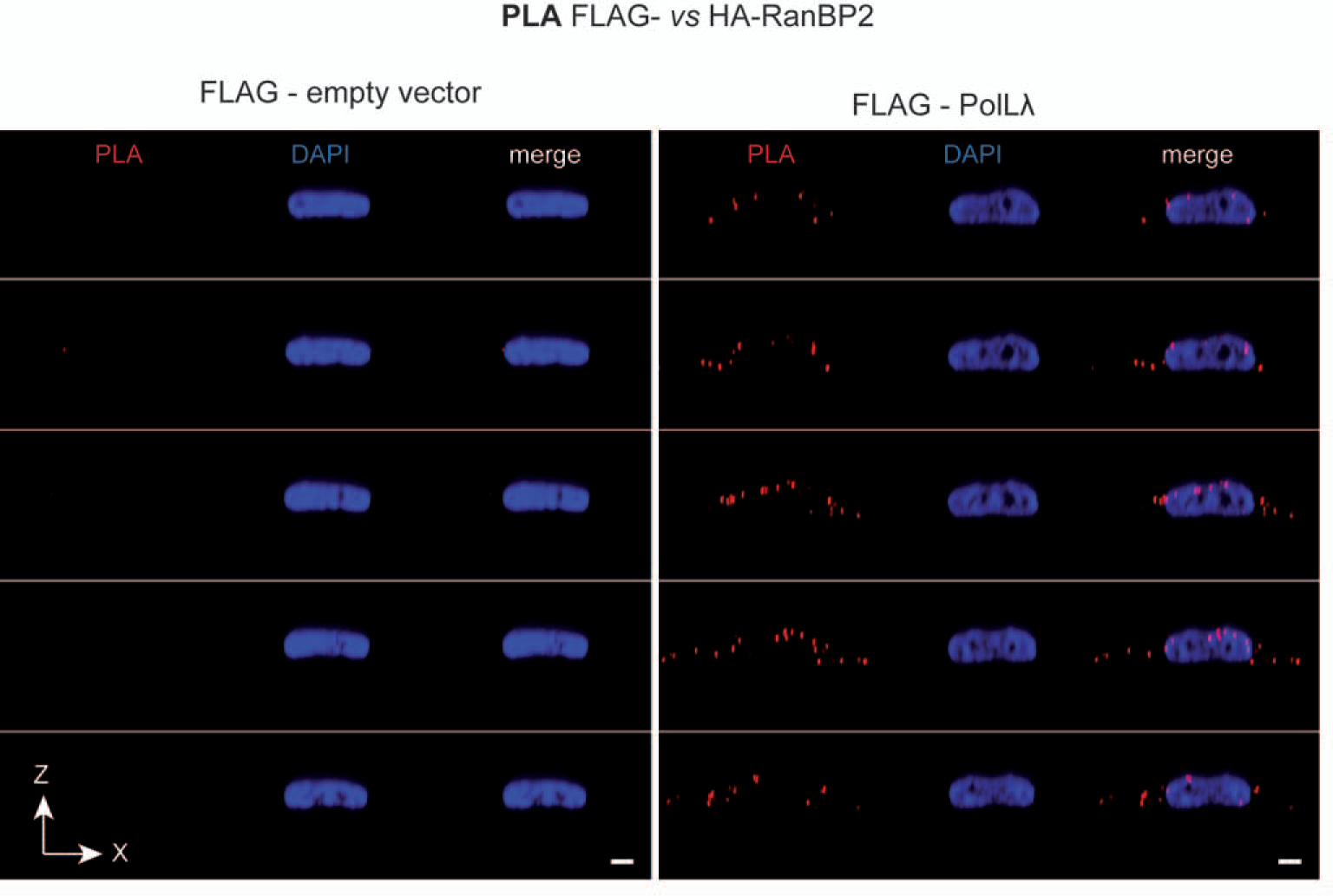
RanBP2-Polλ co-localization at the nuclear pore complex. Human U2OS cells were transiently transfected with a plasmid encoding for HA-tagged RanBP2 together with either Flag-tagged Polλ WT or Flag-empty vector (ev). Proximity ligation assays (PLA) were performed by using mouse monoclonal anti-RanBP2 and rabbit anti-Polλ antibodies and subcellular localization of PLA foci was performed by using confocal laser scanning microscopy. RanBP2-Polλ proximity is indicated with red PLA foci. Nuclei were stained with DAPI (blue). Complete series of X/Z views are shown, with merged images also included in the right. Scale bars, 10 µm.

**Supplemental Figure 7.**
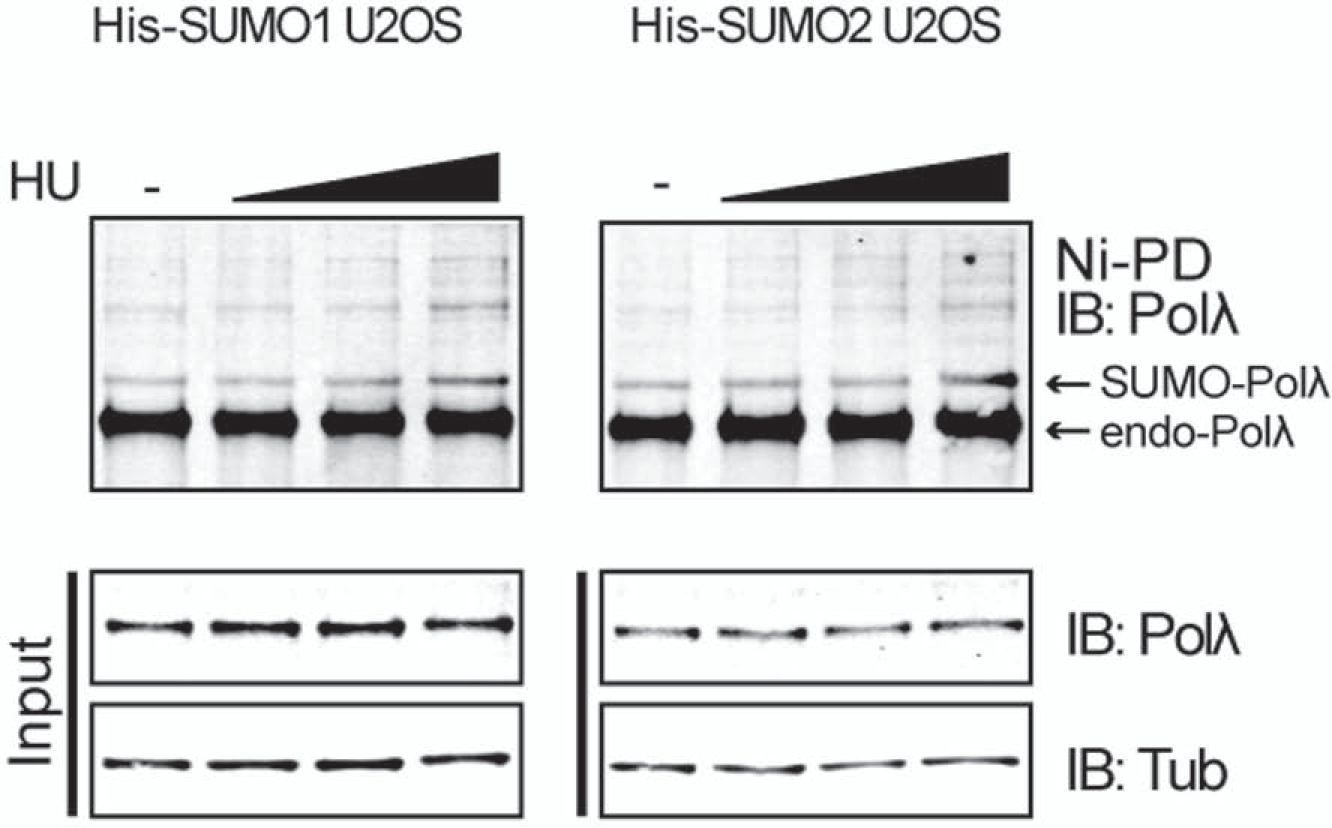
Increased SUMOylation of Polλ in response to HU. His-SUMO1 and -SUMO2 U2OS cells were grown in p100 dishes until 80% confluence, either mock-treated or treated with increasing doses of HU (1 mM, 2 mM and 3 mM) for 2 h, and the harvested. His-SUMO-conjugates were pulled down on Ni-NTA beads (Ni-PD) under denaturing conditions and endogenous SUMO-Polλ was detected by immunoblotting (IB) using anti-Polλ antibody. Expression of Polλ and tubulin was monitored by immunoblotting cell lysates (input) with the corresponding antibodies.

